## appendix for "RNAmigos2: Fast and accurate structure-based RNA virtual screening with semi-supervised graph learning and large-scale docking data"

In this document, we start by providing additional results in A. Then, we analyze our data about binding sites in section B and about our ligand compounds in section B. Finally, we give details about our docking benchmark procedure in section D and other details on our implementation in E

### A Additional Results

#### A.1 Detailed Model Ensembling

In this section, we give our results in the form of numbers, for easier later benchmarking. In Table 1, we provide the results of merging the outputs of two methods, to be compared to its first column where models are used in isolation.

| | Alone | Compat | Aff | rDock | Rnamigos |
|---|---|---|---|---|---|
| Compat | $84.4 \pm 3.4$ | $89.7 \pm 2.9$ | $97.2 \pm 0.9$ | $98.0 \pm 1.2$ | - |
| Aff | $93.9 \pm 0.6$ | - | $94.2 \pm 0.4$ | $96.2 \pm 0.1$ | - |
| rDock | $95.9 \pm 0.0$ | - | - | - | $98.4 \pm 1.1$ |

Table 1: AuROC obtained by different method combinations. All numbers are averaged over three independent seeds and the variation is an estimate of standard variation. In the first column, we do not merge and instead report the performance of models used in isolation. When models are mixed with themselves, it means that we ensemble two instances of the same model. The last cell of the table amounts to the result of the RNAmigos++ model.

It can be seen that different seeds give an overall stable result. Ensembling two Aff does not result in a significant boost. Ensembling two Compat does yield a boost, but only a mix of the two sources of information enables outperforming rDock. Finally, we see that adding rDock results consistently helps, and that merging it with either Mixed or Compat (effectively replacing the docking surrogate with true docking) yields comparable results, with an edge for RNAmigos++.

#### A.2 Fingerprint Prediction

To compare to RNAmigos, we use two ligands sets of decoys for our pockets. The *pdb* set is the set of 264 co-crystallized binders present in the PDB. The *decoyfinder*, consists in non-binders generated for each active using DecoyFinder[10]. For fairness, in this section, we follow the original validation protocol, except for removing ligands based on our filters. We use the *DecoyFinder* and PDB ligands sets as decoys sets and emphasize that at this step, we ignore the ChEMBL compounds. For each pocket in the test set, we perform a virtual screening and aim to retrieve the native binder among decoys. We present the results in Table 2.

| | *DecoyFinder* | PDB Ligands |
|---|---|---|
| RNAmigos | $0.609 \pm 0.294$ | $0.627 \pm 0.284$ |
| fp | $0.821 \pm 0.272$ | $0.856 \pm 0.219$ |
| fp + pre | **0.858± 0.253** | **0.872± 0.210** |

Table 2: AuROC values computed on *DecoyFinder* and PDB decoy sets. We compare those values for the original RNAmigos results and our models in the fp setting, with and without pretraining.

On this more challenging filtered data set, we reproduce the ability of RNAmigos to retrieve a binding affinity signal. Moreover, we see that the proposed enhancements yield a significant performance gain of over 25% AuROC points on the *DecoyFinder* decoy set, achieving a new state-of-the-art on this task. We observe that as expected, pretraining our RNA encoder boosts the performance of our method.

#### A.3 Comparison between pocket and prediction similarity

We compare the similarity of our prediction for two pockets to the ones between these pockets assessed with RMscores. To compare our predictions, we compute a Jaccard score over the set of our top 5% predictions. We order pockets using a 1Dt-sne to find a smooth ordering and display this comparison in figure 6. We can see that there is a group of similar pockets, yet below our 0.7 cutoff, that also results in a group of similar predictions. However, our predictions are similar but not identical for related binding sites, indicating that it is able to be specific.

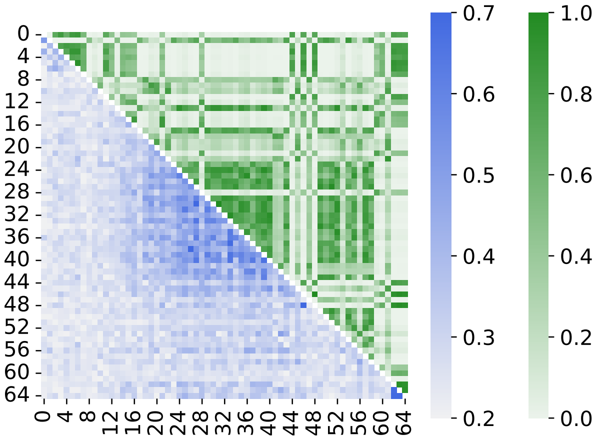


Figure 6: Comparison between RMscores of pockets (lower diagonal) and similarity of predictions (upper one).

#### A.4 Additional ROBIN results

We then turn to assessing the performance of the model on ROBIN. First, we present the performance of different models split by targets, including diversity, AuROCs and enrichment factors, and present results in Table 3. Our results show that rDock and `RNAmigos2` perform similarly, but that their combined usage results in enhanced performances overall. It also shows that our methods overall yield higher diversity albeit slightly reduced enrichment factors.

We then include comparison of the distributions of actives in the dataset, in Figure 7**a**. It shows that the pockets have comparable but different actives. In particular, the ZTP pocket appears to stand out compared to other ones. We also compare actives retrieved by different methods, and show a lower diagonal, which indicates agreement between methods and specificity. However, the diagonal terms are non-zero, which hints at the complementarity of the retrieved solutions by the two methods.

In Figure 8, we show a t-sne of active ROBIN compounds. Additionally, for each pocket, we show the compounds retrieved by our methods as different rows, highlighting which model retrieves which actives. As can be seen in this plot, the AFF model tends to have more grouped predictions of highly populated clusters, while the Compat model tends to have more diverse, scattered and pocket-specific retrieval.

Finally, in Figure 8, we show our performance in the t-sne space that includes inactive compounds, and is shared among pockets. We see that our retrieved actives span a large fraction of the chemical space.

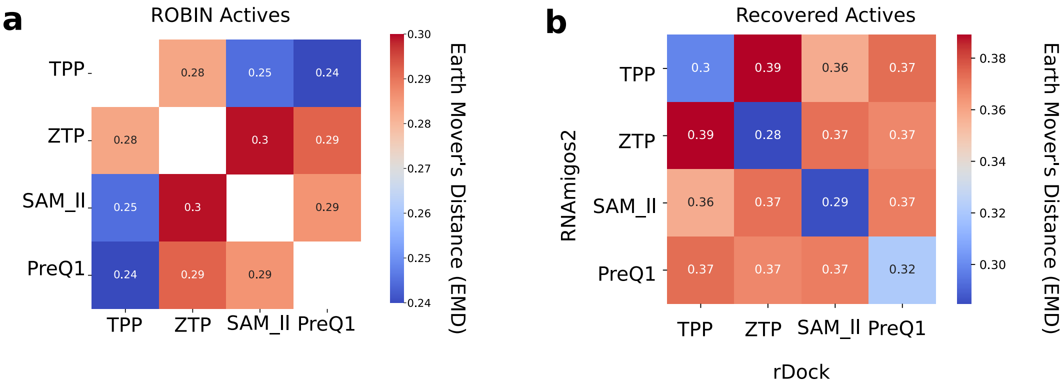


Figure 7: a. Earth Mover's Distance (EMD) between distribution of active compounds for each ROBIN target. b. EMD between *predicted* actives (true positives) from RNAmigos2 and rDock for all pairs of ROBIN targets.

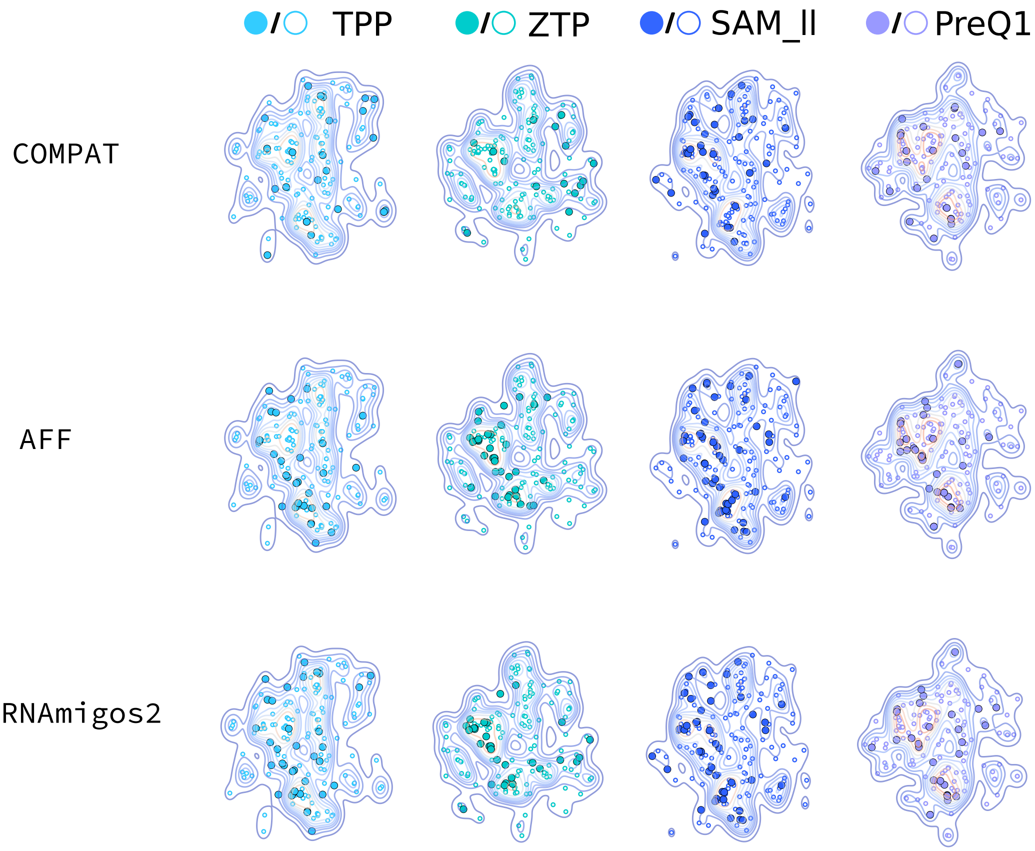


Figure 8: Distribution of ROBIN actives by each sub-model and the combined model `RNAmigos2`. Correctly identified compounds are filled in circles and missed compounds are empty compounds.

| | | AuROC | Diversity | Hopkins | EF1% | EF2% | EF5% |
|---|---|---|---|---|---|---|---|
| PreQ1 | Aff | 0.62 | 0.56 | **0.651** | 1.20 | 1.80 | 2.04 |
| | Compat | 0.59 | 0.53 | 0.626 | 0.60 | 0.60 | 1.44 |
| | RNAmigos2 | 0.62 | 0.58 | 0.641 | 1.80 | 0.90 | 1.68 |
| | RNAmigos++ | **0.65** | **0.59** | 0.648 | 3.59 | 3.29 | 2.40 |
| | rDock | 0.61 | **0.59** | 0.639 | **4.79** | **3.89** | **3.47** |
| SAM_ll | Aff | 0.67 | 0.56 | **0.644** | 5.09 | 3.56 | 3.15 |
| | Compat | 0.61 | 0.58 | 0.628 | 1.53 | 1.52 | 1.63 |
| | RNAmigos2 | 0.66 | 0.59 | 0.643 | 5.09 | 3.30 | 2.64 |
| | RNAmigos++ | **0.68** | **0.60** | 0.642 | 4.58 | **5.08** | **3.45** |
| | rDock | 0.66 | 0.58 | 0.642 | **5.60** | 4.83 | **3.45** |
| TPP | Aff | 0.64 | 0.56 | 0.639 | 3.68 | 3.68 | **2.95** |
| | Compat | 0.56 | 0.56 | **0.658** | 0.00 | 1.53 | 1.47 |
| | RNAmigos2 | 0.62 | **0.58** | 0.652 | 3.68 | 1.84 | 2.33 |
| | RNAmigos++ | **0.65** | **0.58** | 0.643 | 5.52 | **4.30** | 2.58 |
| | rDock | 0.62 | 0.57 | 0.627 | **6.14** | 3.68 | **2.95** |
| ZTP | Aff | **0.66** | 0.58 | 0.673 | 1.17 | 2.34 | **2.93** |
| | Compat | 0.39 | 0.58 | 0.587 | 0.00 | 0.88 | 0.94 |
| | RNAmigos2 | 0.61 | **0.63** | **0.680** | 1.17 | 0.88 | 2.22 |
| | RNAmigos++ | 0.61 | **0.63** | 0.665 | 0.00 | 1.75 | 2.11 |
| | rDock | 0.63 | 0.60 | 0.672 | **1.76** | **4.09** | **2.93** |

Table 3: Model performance on ROBIN.

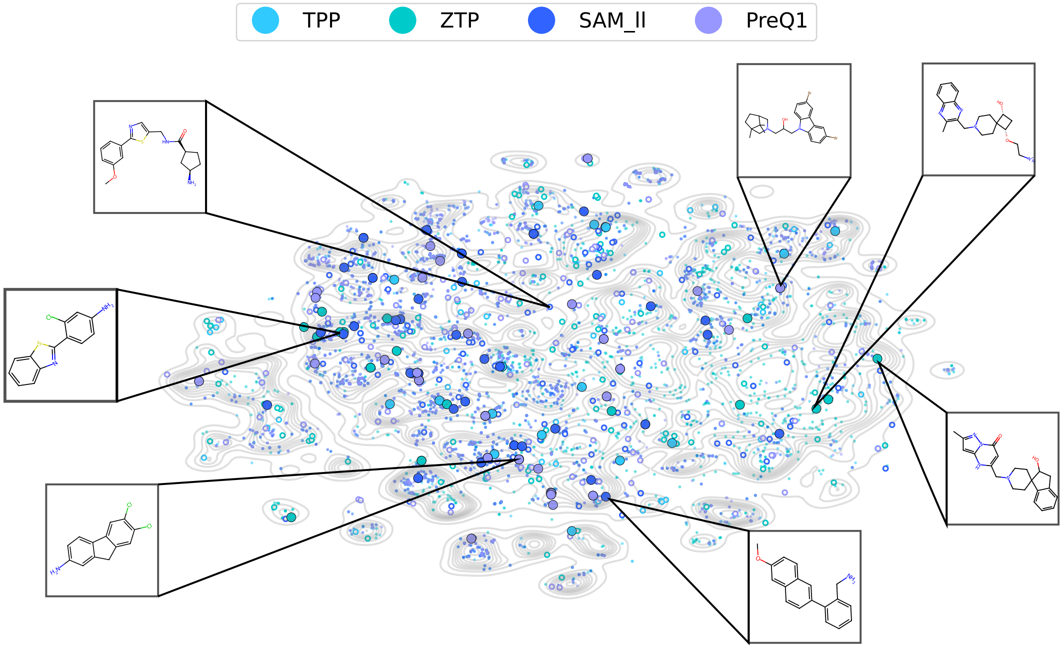


Figure 9: ROBIN chemical space. Each ligand is a point in t-SNE space. Active compounds for each RNA are colored. Filled-in points are actives which RNAmigos2 correctly identifies and hollow points are missed actives. Contour lines represent density of whole chemical space.

### B Properties of RNA binding sites

In this section, we analyze the properties of our binding sites and their representation as 2.5D graphs. First, we assess the redundancy of our binding sites. We use RMScore[80] to measure and analyze RNA structural similarity on our binding sites. The distribution of those values, shown in Figure10a), displays a strong peak below 0.3. We compute a multidimensional scaling [49] of our data based on this similarity measure in Figure 10b. In this representation, dissimilarity of the samples correlates with the distance between points on the plot. The absence of clusters in this representation suggests that the data-set is not redundant. When trying to enforce a clustering using RMscore values, the silhouette score has a value below 0.3, showing that the clusters are bad and hence, that sites are not similar.

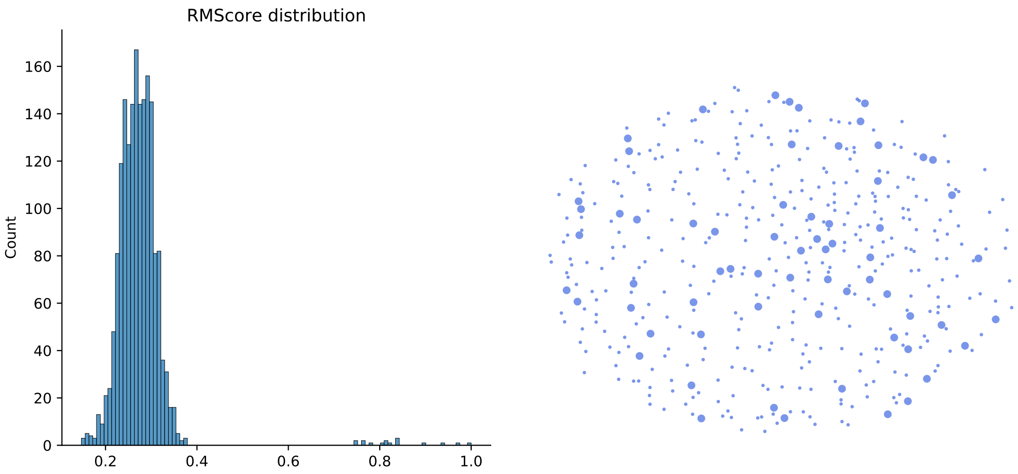


(a) RMScore distribution over binding pockets.

(b) 2D MDS projection of pockets. Pockets in the test sets are shown as larger points

Figure 10: Pocket structural similarity analysis.

We now turn to an analysis of the representation of our binding sites. Table 4 reports the main features of the graphs we extracted for our 1740 binding sites.

| | Nodes | Edges | Non-canonicals | Canonicals | Connected components |
|---|---|---|---|---|---|
| Mean | 10.3 | 11.6 | 1.47 | 10.1 | 1.20 |
| Std | 4.0 | 5.9 | 2.1 | 5.3 | 0.42 |
| Min. | 4 | 2 | 0 | 0 | 1 |
| 25% | 7 | 6 | 0 | 5 | 1 |
| 50% | 11 | 12 | 1 | 10 | 1 |
| 75% | 13 | 16 | 2 | 15 | 1 |
| Max. | 22 | 33 | 18 | 23 | 3 |

Table 4: Descriptive statistics of the binding sites.

### C Properties of ligands

Next, we turn to analyzing the ligands used in this paper. We compare properties of our two ligand sets. As a reminder, our first set is 264 ligands filtered from the PDB, used to compare to RNAmigos and to train our `Compat` model. Our second set is made of 500 ligands per binding site extracted from the ChemBL database and selected for diversity and drug-likeness. We present a few properties of those compounds in Table 5. Moreover, we compute the distribution of inter-molecular energies for the ligands (Figure 11a) and the distribution of pairwise similarities (Figure 11b).

| Descriptor | PDB | ChEMBL |
|---|---|---|
| Molecular weight | 455.5 | 271.9 |
| Logp | -0.083 | 1.315 |
| H bond donor | 4.467 | 1.776 |
| H bond acceptors | 9.05 | 4.6 |
| Rotatable bonds | 6.3 | 2.3 |
| Number of atoms | 31.6 | 19.2 |
| Molar refractivity | 113 | 71.7 |
| Topological surface area mapping | 159 | 74 |
| Formal charge | 0.20 | 0.04 |
| Heavy atoms | 31.6 | 19.3 |
| Number of rings | 3.2 | 2.9 |

Table 5: Chemical descriptors of PDB Ligands and ChEMBL compounds

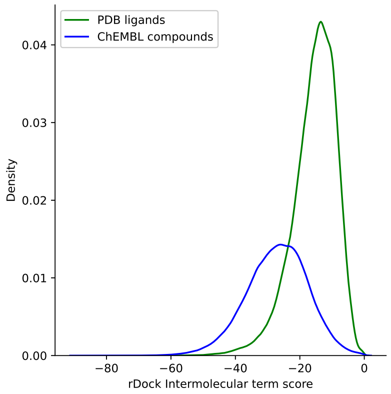


(a) Rdock score distribution for the PDB and ChEMBL datasets.

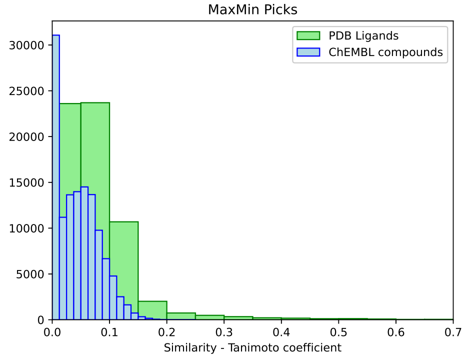


(b) Distribution of pairwise distances for the PDB and ChEMBL datasets.

We can see that the PDB ligands do not display drug-like features. In particular, the compounds are bigger than drug-like compounds. They also display worse docking scores on average. These results indicate that this second ChEMBL was indeed needed for an assessment of our tool from a drug design perspective. Finally, our results show that the samples in each of the data sets are diverse.

Finally, in Figure 12 we plot a 2D MDS of our ligand space in which we identify test compounds. This illustrates that our actives span a large fraction of the chemical space.

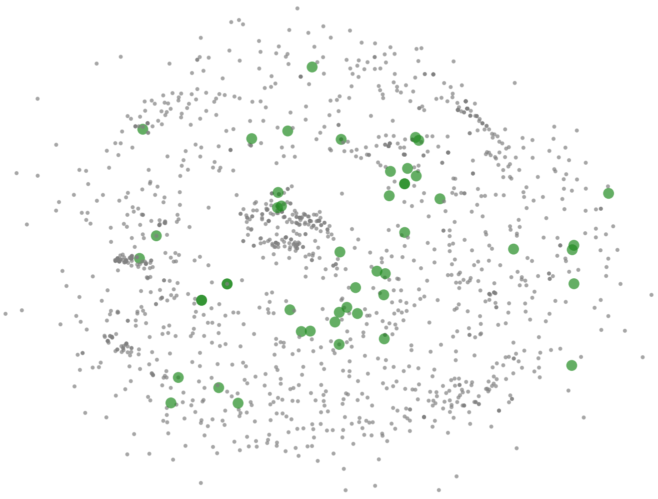

Figure 12: RMScore distribution over binding pockets.

### D Docking software benchmark

We assessed three docking tools, rDock [55], DOCK 6[31] and AutoDock Vina[68] which were identified in a recent review of nucleic-acid docking tools[38]. rDock was developed for NA-ligand docking, DOCK 6 is an extended version of DOCK 5 and specifically optimized for RNA and AutoDock Vina was used without any specific optimization for NAs In addition, we used AnnapuRNA[61], a knowledge-based scoring function, as an independent referee to compare these candidate software in terms of native pose identification. We first establish our docking procedure and compare our candidate docking programs using self and cross-docking experiments. In self-docking experiments, we evaluate programs' ability to predict the binding modes inside the native macromolecule by comparing the generated ligand poses to the crystal structure. In cross-docking experiments, we evaluate the ability of docking tools to retrieve actives among decoys. Based on both experiments, we choose to use rDock as our docking program.

Let us describe the settings with which we have run our docking experiments, using rDock, DOCK 6 and AutoDock Vina. For AutoDock Vina, we defined the exhaustiveness, energy_range and num_modes to be 8, 3 and 100, respectively. The search space (i.e., the coordinates for the center of mass and the size of the binding sites) was calculated by AutoGridFR [79]. We also executed a different set of experiments by using the parameters presented in [55] (i.e., exhaustiveness=16, num_modes=100, energy_range=30 and the search space calculated by rDock). The greatest affinity difference between (the best pose predicted by) the two sets of parameters was 0.3 kcal/mol. Regarding the RMSD score, the greatest difference occurred for the 1BYJ structure (5.404); however, all of the other structures presented differences lower than 0.183. These results agree with the developers of AutoDock Vina who stated that *"With the default (or any given) setting of exhaustiveness, the time spent on the search is already varied heuristically depending on the number of atoms, flexibility, etc. Normally, it does not make sense to spend extra time searching to reduce the probability of not finding the global minimum of the scoring function beyond what is significantly lower than the probability that the minimum is far from the native conformation"*.

Three steps are necessary to run rDock: 1) system definition, 2) cavity generation, and 3) docking. As in [55], the cavity was defined using the crystallographic ligand (found in the PDB data-base) as reference through the "reference ligand method" implemented in rDock. All the parameters were used by default; but the radius, small_sphere, max_cavities, flexibility were set to 4.0, 1.0, 1 and 3, respectively. The models were scored using the "dock_solv.prm" function.

To perform a docking with DOCK 6, we used the parameters by default and followed the steps to prepare the receptor and ligand structures, generate the molecular surface of the receptor, generate the spheres surrounding the receptor, select a subset of spheres to represent the binding site using the crystallographic ligand, to create the grid used to scoring and finally to run the flexible ligand docking experiments.

Self and cross docking experiments are performed to have a fair comparison between the capabilities of the selected tools when used on RNA molecules as a target.

#### D.1 Self-docking experiments

During the self-docking experiments, we evaluate the ability of the docking tools to predict the binding modes inside the native macromolecule. Therefore, we compare the generated ligand poses predicted as best to the crystal structure. We use the root mean square deviation (RMSD) as the metric to determine the accuracy of the prediction.

The experiments were run on a set of 9 RNA-ligand complexes (1EI2, 1QD3, 1LVJ, 1BYJ, 1NEM, 1PBR, 1TOB, 2TOB, and 1FMN) obtained from [6] and [79]. For DOCK 6 and AutoDock Vina, the complex structures were downloaded from the PDB data-base. For rDock, the structures were obtained from the validation dataset reported by [55].

Table 6 summarizes the descriptive statistics for the RMSD of the 20 best poses for each complex and each tool. The total number of RSMD values for each docking tool is 180 (20 for each complex). The mean and median RMSD are more favourable for rDock with 4.28A and 3.62A, respectively (6.17 and 6.67 for DOCK 6 and 7.21A and 7.43 for AutoDock Vina).

| Docking tool | Mean | Std. | Min. | 25% | 50% | 75% | Max. |
|---|---|---|---|---|---|---|---|
| rDock | 4.28 | 2.85 | 0.68 | 1.84 | 3.62 | 6.78 | 10.32 |
| DOCK 6 | 6.17 | 4.45 | 0.88 | 1.92 | 6.67 | 7.96 | 14.85 |
| AutoDock Vina | 7.21 | 2.02 | 1.17 | 5.79 | 7.43 | 8.84 | 10.74 |

Table 6: Descriptive statistics of the RMSD obtained with rDock, DOCK 6 and AutoDock Vina

Based on the self-docking experiment, we can conclude that rDock has better performance than DOCK 6 and AutoDock Vina in docking ligands against RNA targets.

#### D.2 Cross-docking experiments

In cross docking experiments, we evaluate the ability of docking tools to retrieve actives among decoys. We do so by docking each ligand in the dataset to each RNA target. The experiments were carried out on a set of 50 RNA-ligand complexes, having 31 unique ligands. We use the native ligand of each complex as the active ligand and the other 30 ligands in the dataset as inactive ligands.

AutoDock Vina only returns the binding affinity of the bound conformations calculated with its scoring function, DOCK 6 returns the grid score (sum of the van der Waals and electrostatic interactions) and rDOCK, in addition to the total score, details the value of each term of the scoring function (intermolecular, ligand intramolecular, site intramolecular and external restraint). Beside the total score, we also ranked the poses predicted with rDock based on the intermolecular term because it represents the RNA-ligand interaction score [55]. We use AnnapuRNA to re-score the poses. The results are presented in Table 7 each docking tool and for the poses. We find rDock with the intermolecular term of its scoring function to have the best performance. Based on the results shown in this section, we chose rDock as our docking tool.

| | rDock Total | rDock Inter. | DOCK 6 | Vina | rDock AnnapuRNA | Vina AnnapuRNA | DOCK6 AnnapuRNA |
|---|---|---|---|---|---|---|---|
| AuROC | 0.68 | 0.72 | 0.61 | 0.66 | 0.60 | 0.62 | 0.53 |

Table 7: Mean of the AuROC scores obtained with different docking software. We compare the use of the Rdock score, the use of only its intermolecular term, Rdock with Annapurna, DOCK 6, DOCK 6 with AnnapuRNA and AutoDock Vina with Annapurna.

#### D.3 Runtime for docking experiments

Table 8 reports the execution time it took to perform the docking experiments, including the cavity generation stage and the docking itself. The method selected for generating the cavity was to take the native ligand as the reference ligand.

| Cavity generation | | Docking - ChEMBL | Docking - PDB |
|---|---|---|---|
| # Pockets | Time (minutes) | Avg time per compound (min.) | Avg time per compound (min.) |
| 645 | 5 | | |
| 85 | 8 | | |
| 335 | 20 | | |
| 302 | 40 | 0.41 | 1.5 |
| 210 | 60 | | |
| 35 | 120 | | |

Table 8: Docking experiments runtimes.

The approach rDock uses to parallelize the jobs is independently running each molecule, splitting the ligands file into multiple files, and running each independently. To parallelize the docking experiments, we executed the experiments over each pocket in parallel using an array job in Compute Canada. Therefore, for the cavity generation task we used as many CPU numbers as pockets (1740), taking approximately 2 hours to complete this task. In the same sense, the docking of each pocket with the ChEMBL compounds, using 1740 CPUs, is approximately 4 hours and for the PDB ligands, 7 hours.

### E Additional details of the methods

#### E.1 Model Architecture and Training

The RNA encoder $f_\theta$ is implemented as a Relational Graph Convolutional Network (RGCN) [58] with 3 convolutional layers, and a hidden dimension of 64 and no basis sharing. The pretraining of this model is done using the RNAGlib package [40] with default parameters. The resulting node embeddings are pooled into a graph-level embedding $\phi_\mathcal{G}$ with a global attention pooling layer. The OptiMol ligand encoder is a 3 layer GCN with 56 hidden dimensions. The decoder heads which take the concatenated RNA and ligand embeddings is a 3-layer multi-layer perceptron with 32 hidden dimensions at each internal layer. We use a dropout probability of 0.2 and apply batch normalization at each layer. Encoder models and data loaders are implemented in DGL [71] and training is done in PyTorch [48].

We train our networks using the Adam optimizer for 1000 epochs `Compat` setting, and for 20 epochs in the `Aff` setting.

#### E.2 Perturb algorithms

We provide a detailed algorithm for how we compute the perturbed version of our pockets in the following algorithms. We use algorithm 3 with the two sampling strategies described respectively in algorithms 1 and 2.

**Algorithm 1:** Noised sampling algorithm.

**Data:**

- A pocket: $p$
- A number of nodes $n$
- A whole RNA graph $R$

**Result:** A set of nodes of $R$

$S \leftarrow$ `random_choice`$(p, n)$ ;
**return** $S$

**Algorithm 2:** Shifted sampling algorithm.

**Data:**

- A pocket: $p$
- A number of nodes $n$
- A whole RNA graph $R$

**Result:** A set of nodes of $R$

border $\leftarrow$ `get_border`$(p)$;
seed $\leftarrow$ `sample`(border);
prev_out $\leftarrow \emptyset$;
out $\leftarrow$ {seed};
**while** $|out| \leq n$ **do**
    prev_out $\leftarrow$ out;
    out $\leftarrow$ `bfs`(out, 1, $R$);
**end**
missing $\leftarrow n -$ |prev_out|;
border $\leftarrow$ out $-$ prev_out;
last $\leftarrow$ `random_choice`(border, missing);
$S \leftarrow$ last $\cup$ prev_out;
**return** $S$

**Algorithm 3:** Perturbation algorithm. Perturb a dataset of pockets with a certain sampling algorithm.

**Data:**

- Set of pockets: $\mathcal{P}$
- Sampling algorithm: `sample`
- Set of size fractions: **F**
- Set of hops: **H**
- Number of replicates: $I$

**Result:** A set of perturbed pockets $\tilde{\mathcal{P}}$

$\tilde{\mathcal{P}} \leftarrow \emptyset$ **for** $p$ *in* $\mathcal{P}$ **do**
    $R \leftarrow$ `get_rna`$(p)$;
    **for** $h$ *in* **H** **do**
        $p_h \leftarrow$ `bfs`$(p, h, R)$;
        **for** $f$ *in* **F** **do**
            $n_f \leftarrow$ int$(f|p|)$;
            **for** $i$ *in* $I$ **do**
                $\tilde{p} \leftarrow$ `sample`$(p_h, n_f, R)$;
                $\tilde{\mathcal{P}} \leftarrow \tilde{\mathcal{P}} \cup \tilde{p}$;
            **end**
        **end**
    **end**
**end**
**return** $\tilde{\mathcal{P}}$
